# Neural networks implicated in autobiographical memory training

**DOI:** 10.1101/2021.10.05.463255

**Authors:** Dragoş Cȋrneci, Mihaela Onu, Claudiu C. Papasteri, Dana Georgescu, Catalina Poalelungi, Alexandra Sofonea, Nicoleta Puşcaşu, Dumitru Tanase, Teofila Rădeanu, Maria-Yaelle Toader, Andreea L. Dogaru, Ioana R. Podină, Alexandru I. Berceanu, Ioana Carcea

## Abstract

Training of autobiographical memory has been proposed as intervention to improve cognitive functions. The neural substrates for such improvements are poorly understood. Several brain networks have been previously linked to autobiographical recollections, including the default mode network (DMN) and the sensorimotor network. Here we tested the hypothesis that different neural networks support distinct aspects of memory improvement in response to training on a group of 59 subjects. We found that memory training increases DMN connectivity, and this associates with improved recollection of cue-specific memories. On the contrary, training decreased connectivity in the sensorimotor network, a decrease that correlated with improved ability for voluntary recall. Moreover, only decreased sensorimotor connectivity associated with training-induced decrease in the TNFα immunological factor, which has been previously linked to improved cognitive performance. We identified functional and biochemical factors that associate with distinct memory processes improved by autobiographical training. Pathways which connect autobiographical memory to both high level cognition and somatic physiology are discussed.

## Introduction

Autobiographical memories represent the fabric of our identity. In several neurodegenerative disorders that manifest with memory loss, self-identity dissipates, inducing anxiety, confusion and impaired social interactions. Training autobiographical recall might be a beneficial behavioral intervention that preserves most defining memories. Several studies have investigated the effects of autobiographical memory training on emotional well-being (Kohler, et al., 2015; Hitchcock, et al., 2016; Serrano, et al., 2004; Ricarte, et al., 2012). However, unlike studies on training of working memory or of spatial memory (de Marco et al., 2016; Brinke et al., 2017), very little is known about how autobiographical training changes the ability to recall autobiographical events, and what neurophysiological substrates might be engaged by training.

Previous imagistic studies have shown that brain regions activated during autobiographical recall form a network that largely overlaps with the default mode network (DMN), comprised of medial prefrontal cortex, medial parietal cortex, lateral and medial temporal lobe, precuneus, posterior cingulate cortex, retrosplenial cortex and temporo-parietal junction (Buchanan, 2007; Spreng et al., 2009). The structures are also activated by imagining future events, navigation and theory of mind, mental processes that require *scene construction* (Hassabis et al., 2007; Hassabis and Maguire, 2007; Spreng et al. 2009). *Scene Construction Theory* (Mullally and Maguire, 2013; Clark, et al., 2020) views the retrieval of episodic memories as a re-constructive process common to several other cognitive functions. According to this theory, we expect that training autobiographical memory retrieval would lead to changes in the activity and connectivity of DMN structures.

A different theory for autobiographical memory recall (and more broadly for episodic memory) is that of *embodied memory*, where recalls rely on sensorimotor simulations of events (Iani 2019). This theory proposes that the patterns of brain activity required during memory encoding will be reactivated at memory recall (Nyberg, et al., 2001; Nyberg, 2002; Iani, 2019). This theory is supported by imaging studies that find activation of sensory and motor areas during episodic memory recall (Nilsson, et al., 2000; Nyberg, et al., 2001; Masumoto, et al., 2006; Mineo, et al., 2018). Based on this theory, we predict that autobiographical training changes activity patterns and/or connectivity within sensorimotor networks.

In addition to changes in neural activity, improvements in memory could also associate with biochemical changes. Autobiographical recalls have been shown to decrease the levels of tumor necrosis factor alpha (TNFα), of interleukin-2 and of interferon gamma (Matsunaga, et al., 2011; Matsunaga, et al., 2013). The relationship between inflammatory state and neural network activity and connectivity is complex. At baseline, blood levels of cytokines associate with changes in the activity of the DMN, limbic, ventral attention and corticostriatal networks, and with changes in connectivity within the DMN (Kraynak, et al., 2018; Marsland, et al., 2017). In relation to autobiographical recall, whereas for interferon gamma an anti-correlation was found with activation of the orbitofrontal and posterior cingulate cortex (Matsunaga, et al., 2013), for TNFα it remains to be determined if such a functional association exists. Increased levels of TNFα associate with poor cognitive performance and aggravated Alzheimer’s dementia (Hennessy, et al., 2017). It is therefore important to determine if autobiographical training could be beneficial by decreasing cytokine levels.

Our hypothesis is that autobiographical memory training increases efficiency within brain networks involved in memory retrieval. To test this hypothesis, we used olfactory cues to induce recall of autobiographical memories. The choice of olfactory modality was dictated by unpublished data from our lab and also by previous findings describing the efficiency of odor-evoked memories (Matsunaga, et al., 2013; Larsson, et al., 2014; Herz, 2016). Odor-evoked autobiographical recall is a technique used in theater training for student actors, to gain access to personal memories, an exercise inspired by the view on acting and memory of Method acting (Stanislavsky, 2010; Cohen, 2010). To investigate changes in brain activity following autobiographical training that could explain lasting changes in memory performance, we performed resting-state functional MRI scanning at the beginning and at the end of training. We focused on the connectivity within functional networks. To determine if autobiographical training can also change the levels of TNFα, we collected blood samples at the beginning and end of training. We then tested a possible association between cytokine and brain activity dynamics.

Our findings bring important scientific evidence to the translational use of a technique primarily employed in theatrical training. We argue that training of autobiographical memories could be used in therapy for the prevention and treatment of memory loss.

## Methods

All methods and experiments have been approved by The Ethics Committee of National University for Theatre and Film I.L Caragiale Bucharest, and followed the guidelines of the Declaration of Helsinki. All participants provided written informed consent for their participation. Subjects: An experimental group of 29 subjects (25 women and 4 men) with a mean age of 34.6 years and a control group of 30 subjects (24 women and 6 men) with a mean age of 32.5 years. Exclusion criteria: rhinitis (or other medical problems that lead to impaired smell), depression, anxiety, chronic diseases that cause infection / inflammation, eyeglasses, metal implants, cardiac pacemaker, claustrophobia.

### Materials

15 odors were used: coffee, vinegar, vanilla, cocoa, wine, onion, fresh apples, cinnamon, orange, sanitary alcohol, paint, tobacco, diesel oil, jasmine fragrance and chamomile. The odors were selected and adapted from the stimuli used in previous studies (Chu and Downes, 2002; Gardner, et al., 2012). The odors were presented individually from small containers with perforated lid. Two questionnaires measuring retrieved memories quality and changes in subjective memory recall process were translated and used (Addis, et al. 2004). A first questionnaire containing 3 seven point Likert scales measured retrieved memories’ quality. The first scale asks them about the valence of that specific memory (where 1 is “very unpleasant” and 7 is “very pleasant”), the second one asks them about the vividness of that memory (where 1 is “very faded” and 7 is “very vivid”) and the third one about the personal relevance of that memory (where 1 is “totally unimportant” and 7 is “very relevant”). A second questionnaire containing 3 seven point Likert scales was used for measuring subjective effect upon memory after one month of training. One scale asks to what extent did the subject noticed the onset of spontaneous memories during the day (outside of the experiment) (where1 means “none” and 7 “to a very large extent”). The second scale asks if the subject noticed a greater ease of voluntarily accessing memories, (where 1 means “none” and 7 means “very easy”). The third scale asks to what extent the subject noticed changes in the ease of remembering her/his dreams (where 1 means “nothing” and 7 means “to a very large extent”).

### Procedure

1. Subject inclusion criteria. A hemoleucogram and C-reactive protein (CRP) measurement were used to check for the presence of an infection / inflammation. Only subjects without signs of infection or inflammation were included. From the same blood samples collected from them, the TNF-α levels from lymphocytes has been measured with a high sensitivity ELISA kit.
2. Pre-training session. All the subjects have been exposed to an odor-triggered retrieval session and the subjects have been video monitored during the procedure. After each retrieved memory, a questionnaire has been completed regarding the quality of that memory according to the criteria of valence, vividness and personal relevance. The procedure took 30 minutes. After this session, all the subjects have been scanned using resting-state functional connectivity fMRI procedure.
3. Training session. After the Pre-training session, each subject from the experimental group underwent an autobiographical reminder training for one hour, 2 times / week, for 4 weeks. The experimental group was stimulated to voluntary recall autobiographic memories using 15 odors. Subjects were encouraged to detail the memories as much as possible. After each reminder, a questionnaire was completed aiming at the quality of the reminder according to the criteria of valence, vividness and personal relevance. After the Pre-training session, each subject from the control group watched 2 short movies for 45 minutes, 2 times / week, for 4 weeks. After watching each movie, the subjects evaluated that movie in terms of valence, intensity and dominance.
4. Post-training session. After 4 weeks, all subjects have been exposed to the following assessments: A hemoleukogram and C-reactive protein (CRP) measurement in order to check for the presence of an infection / inflammation, and also for the serum level of TNF-α. All subjects have been exposed to an odor-evoked autobiographical memory recall session, and during this session they have been video monitored. After each retrieved memory, a questionnaire will be completed aiming at the quality of the reminder according to the criteria of valence, vividness and personal relevance. In addition they completed 3 Likert scales regarding the changes they observed after one month of training (ease of voluntarily accessing memories, the onset of spontaneous memories during the day, and the ease of remembering her/his dreams). The procedure took 30 minutes. After this session, all subjects have been scanned using resting-state functional MRI procedure.

We compared the number of odor-evoked memories between Pre-training and Post-training sessions, the scores of valence, vividness and personal relevance between Pre-training and Post-training sessions, the scores of Post-training Voluntary memory scale, the level of TNF-α between Pre-training and Post-training sessions, and changes in brain networks connectivity, in both experimental and control groups.

### Imaging

A 3T Siemens Skyra-MR scanner was used to acquire a resting state functional acquisitions with 281 axial volumes, by means of a 2-dimensional multi-slice echo-planar imaging sequence (TR=2500 ms, TE=30ms, matrix=94×94, voxel size=4×4×4.3mm). Each functional acquisition duration was 11min42s. Additionally, anatomical images were acquired (T1-weighted MP-RAGE, TR/TE=2200/2.51 ms, voxel size 0.9×0.9×0.9 mm). A high-pass temporal filtering cut-off of 100s was applied. The first 5 volumes, acquired to allow longitudinal magnetization to reach a steady state, were discarded.

Data analysis was performed using FMRIB Software Library (FSL) package (http://fsl.fmrib.ox.ac.uk/fsl/fslwiki/). Head motion in the fMRI data was corrected using multi-resolution rigid body co-registration of volumes, as implemented in the MCFLIRT software. For one experimental and one control subjects, the movement was too substantial to be corrected, and data from these subjects was excluded from the rest of the analysis. Brain image extraction was carried out for motion corrected BOLD volumes with optimization of the deforming smooth surface model, as implemented in the BET software. Rigid body registration as implemented in the FLIRT software was used to co-register fMRI volumes to T1-MPRAGE (brain-extracted) volumes of the corresponding subjects and subsequently, to the MNI152 standard space. The images were smoothed with a 5 mm filter.

### Resting state acquisition

Independent Component Analysis (ICA) - the Multivariate Exploratory Linear Decomposition into Independent Components (MELODIC) tool was used to perform spatial group-ICA using multisession temporal concatenation to produce 50 independent component maps (IC maps) representing average resting state networks.

Resting-state networks were identified by visual inspection. The IC maps associated with motion or which were localized primarily in the white matter or CSF spaces were classified using criteria suggested by Kelly et al. (2010) and excluded from further study. We also took into account ICA prominent low-frequency power of Fast Fourier Transformation (FFT) spectra and slow fluctuation in time courses. The remaining 15 networks were identified as classical RSNs as previously reported (Smith et al., 2009; Zuo et al., 2010). The Juelich histological atlas and Harvard-Oxford cortical and subcortical atlases (Harvard Center or Morphometric Analysis) were used to identify the anatomical location, and NeuroSynth 100 top terms atlas (http://neurosynth.org) was used to identify the functional components of the resulting ICA maps.

An intra-network connectivity analysis was performed. This analysis involves comparing the subject-specific spatial maps between experimental and control conditions. To determine subject-specific spatial maps, dual regression analysis was performed on the obtained neural networks using variance normalization (with variance normalization the dual regression reflects differences in both activity and spatial spread of the resting-state networks), similar to previous studies (Emerson et al., 2016; Onu et al., 2015). For the statistical analysis, i.e. the paired two-group difference (two-sample paired t-test), the different component maps were collected across subjects into single 4D files (1 per original ICA map) and tested voxel-wise by nonparametric permutation using the FSL randomize tool (https://fsl.fmrib.ox.ac.uk/fsl/fslwiki/Randomise) with 5000 permutations and a threshold-free cluster enhanced (TFCE) technique to control for multiple comparisons. As we tested a multitude of resting state networks, we addressed the issue of multiple testing correction by controlling the false discovery rate (FDR) at p<0.05.

As we were interested in knowing whether the connectivity values correlate with behavioral parameters as effect of the training program, we further extracted averaged numerical values from the stage 2 maps of the main analysis, for each individual, for the specific clusters where connectivity changes occurred between training and control conditions. The connectivity quantified indices calculated as a result of these procedures were then analyzed in GraphPad Prism.

### Biochemistry

TNFα levels were measured from lymphocytes. Blood samples were obtained by venipuncture using EDTA-coated tubes. 2.5 ml fasting venous blood were used to obtain lymphocytes, which were separated by density gradient centrifugation (Biocoll separating solution, Biochrom GmbH). After separation, the lymphocytes were resuspended in 1ml RPMI culture media (Biochrom GmbH) and ultrasonicated. The supernatant was then aliquoted and stored at -20ºC. Due to technical problems, many of the stored probes were compromised. We were able to use PRE and POST probes from 9 subjects that underwent training and from 5 subjects in the control group. TNFα was measured in these samples using a high sensitivity ELISA kit (IBL International GmbH) with the detection limit of 0.13 pg/ml. The calculated intra-assay coefficient of variation was 8.5% and the inter-assay coefficient of variation was 9.8%. TNFα concentrations were measured using the Tecan Reader, with Magellan Reader software (Tecan Group, Ltd, Switzerland). For the calculation of results we used a 4-parameter curve.

## Results

To determine how training might improve autobiographical memories, we evaluated several aspects related to autobiographical recollections in 29 subjects that underwent 4 weeks of training and in 30 control subjects. The autobiographical memories were triggered with olfactory cues (orange, coffee, etc.) that were presented to experimental subjects in small vials, each at a time. At baseline (Experimental Day 1, PRE) and at the end of the experiment (Experimental Day 10, POST), subjects were presented the cues and asked to recollect an episode from their own life (**Fig. 1A**). In POST, they were asked to score on an analog scale if they observed a change since the start of the experiment in several mnemonic aspects outside of laboratory settings: voluntary recollections, spontaneous recollections or dreams. During the eight days of training, the experimental subjects came to the lab and underwent a similar procedure, where they were asked to recollect autobiographical memories in response to the olfactory cues presented. Control subjects visited the lab the same amount of time, but instead of autobiographical training, they were asked to watch and score a series of videoclips, a control activity meant to match the level of engagement of the experimental group. PRE and POST intervention, both experimental and control subjects were scanned at resting-state for 11.7 min in order to investigate changes in neural network activity induced by training (**Fig. 1A**).

**Figure 1:**
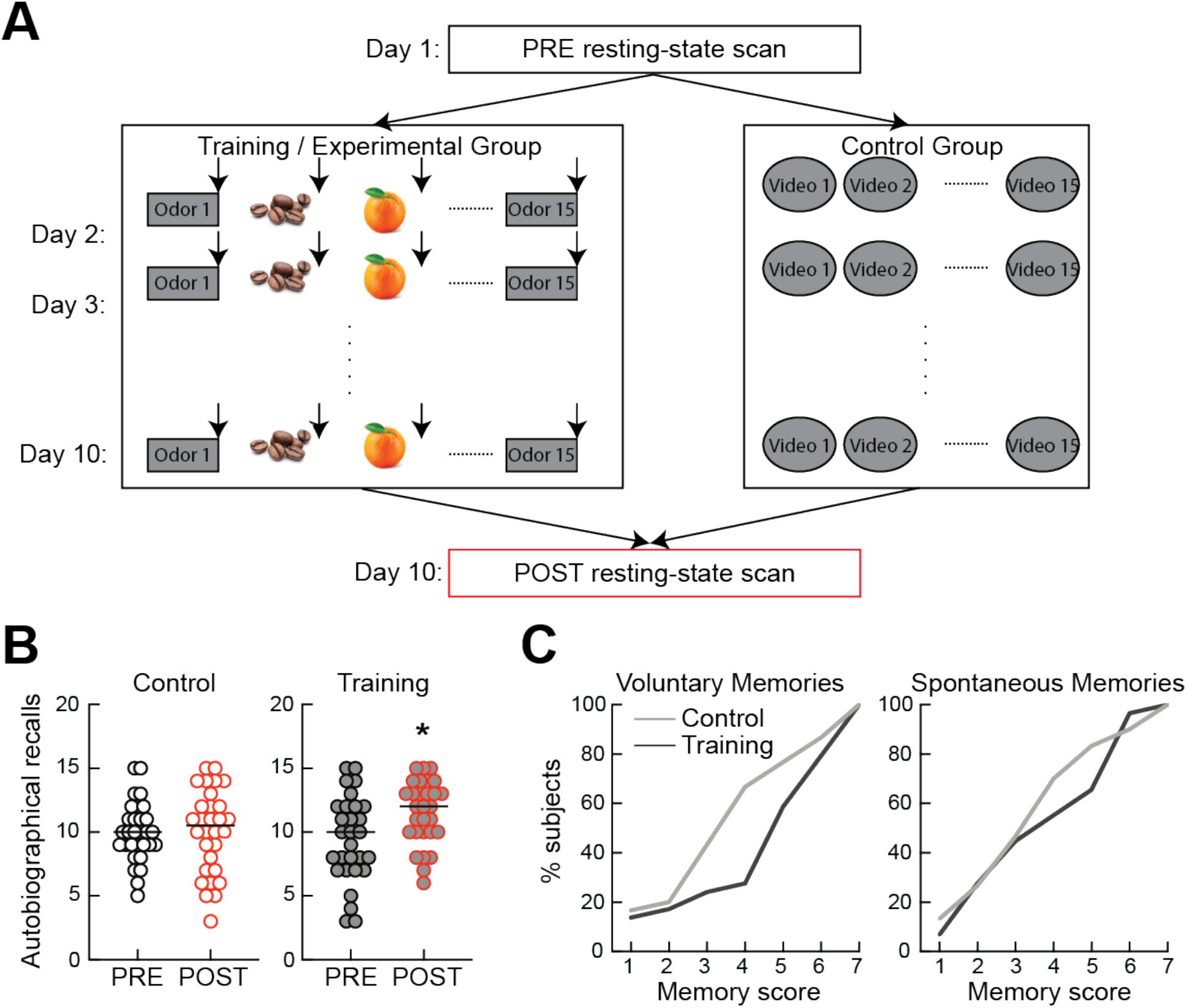
Behavioral effects of autobiographical training. (A) Diagram of the experimental design. (B) Training increases the number of odor-evoked recalls (p=0.0006, N=29). (C) Training improves voluntary recalls in a significant proportion of subjects (p=0.02, N=29). *, p<0.05.

At the behavioral level, we observed that autobiographical training increases the number of odor evoked recalls (**Fig. 1B;** training PRE: 9.4±0.6 recalls/session, training POST: 11.5±0.4, p=0.0006, Wilcoxon’s matched-pairs signed rank test, N=29). The control intervention did not change the number of cue-triggered recalls (**Fig. 1B**; control PRE: 10±0.4 recalls/session, control POST: 10±0.6, p=0.7, N=30). Autobiographical training also improves the ability to recall voluntary memories outside of the laboratory setting. In the control group, only 33.4% of subjects reported an improvement of voluntary memory (score above 4, on a scale from 1 to 7), whereas in the training group 72.4% subjects reported improved voluntary recollection (**Fig. 1C**; p=0.02, Kolmogorov-Smirnov test). The score for spontaneous memories was not affected by training (training: 44.8% subjects reported improved spontaneous memories, control: 50%, p=0.7). Similarly, we did not observe a change in dreams following training (p=0.9, data not shown). Also, training did not change the vividness, valance and personal relevance of autobiographical recalls.

### Autobiographical training increases default mode network connectivity

To investigate whether changes in brain connectivity associate with changes in memory observed after autobiographical training, we acquired resting-state BOLD activity in PRE and POST in all experimental and control subjects. We considered 15 networks, and determined whether training but not control condition change connectivity for either of these networks in POST. We found an increase in the connectivity between the anterior part of the DMN and a region in the right medio-dorsal thalamus (**Fig. 2A,B**; ‘PRE control’ connectivity: -0.07±0.2, ‘PRE training’ connectivity: 0.19±0.14, ‘POST control’ connectivity: -0.44±0.2, ‘POST training’ connectivity: 0.55±0.16; two-way ANOVA, effect of training p=0.002, interaction between time and training p=0.02; Sidak’s multiple comparison correction shows significant difference in POST between training and control, p=0.0003). The strength of DMN-thalamus connectivity positively correlates with the number of recalls during the corresponding autobiographical session (**Fig. 2C**; r:0.3, p<0.001), indicating that it could serve as a mechanism for improved odor-evoked autobiographical memory retrieval. The change in DMN-thalamus connectivity did not correlate with the increase in voluntary memory after training.

**Figure 2:**
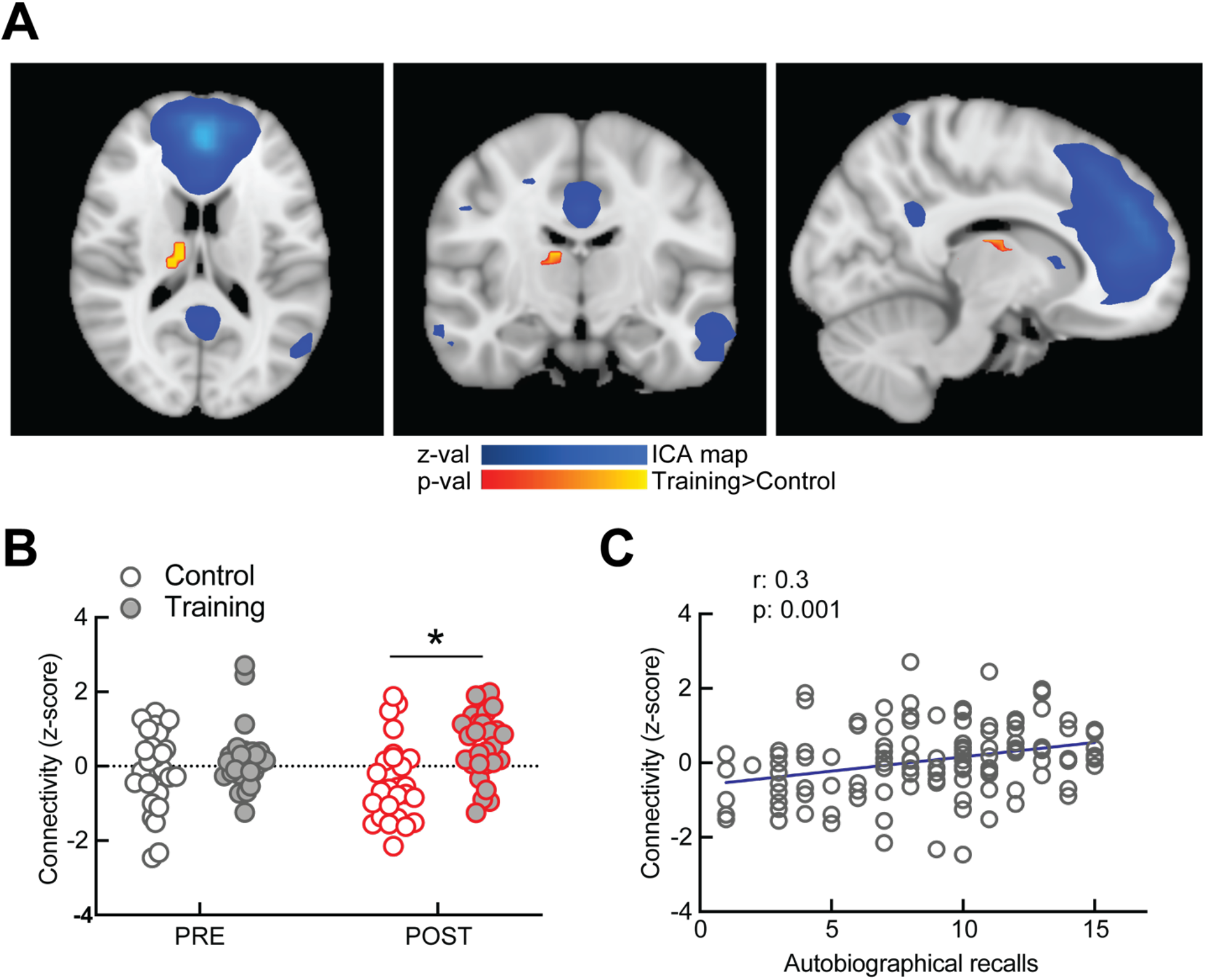
Increased resting state connectivity of anterior DMN after autobiographical training. (A) Horizontal, coronal and sagittal sections showing the part of right mediodorsal thalamus (yellow/red) with increased connectivity with anterior DMN (blue) after training. (B) Summary data showing increased connectivity between right thalamus and anterior DMN after training compared to controls (p=0.0003). (C) Positive correlation between resting-state connectivity and number of odor-evoked recalls. *, p<0.05.

### Autobiographical training decreases connectivity in the sensorimotor network

To understand what patterns of connectivity might explain the increase in voluntary recall after training, we performed voxel-wise correlations with behavioral scores, for all subjects, in POST-training condition. Connectivity within the sensorimotor network was the only one significantly correlated with voluntary memory score after correction for multiple tests. More exactly, connectivity of clusters within the Justapositional Lobule Cortex (formerly Supplementary Motor Cortex) were negatively correlated with voluntary memory score (**Fig. 3A**). We extracted these clusters and went back to subjects-specific connectivity maps (for PRE and POST conditions) and further calculate mean z-scores for these. Intra-network connectivity for the Justapositional Lobule Cortex, part of the sensorimotor network, decreased after the training procedure (**Fig. 3B;** ‘PRE training’ connectivity: 0.50±0.38, ‘POST training’ connectivity: -0.45±0.37, paired t-test, p=0.01), but not after the control intervention (‘PRE control’ connectivity: 0.67±0.49, ‘POST control’ connectivity: 0.70±0.35, p=0.9). There were also no significant differences in PRE between training and control groups.

**Figure 3:**
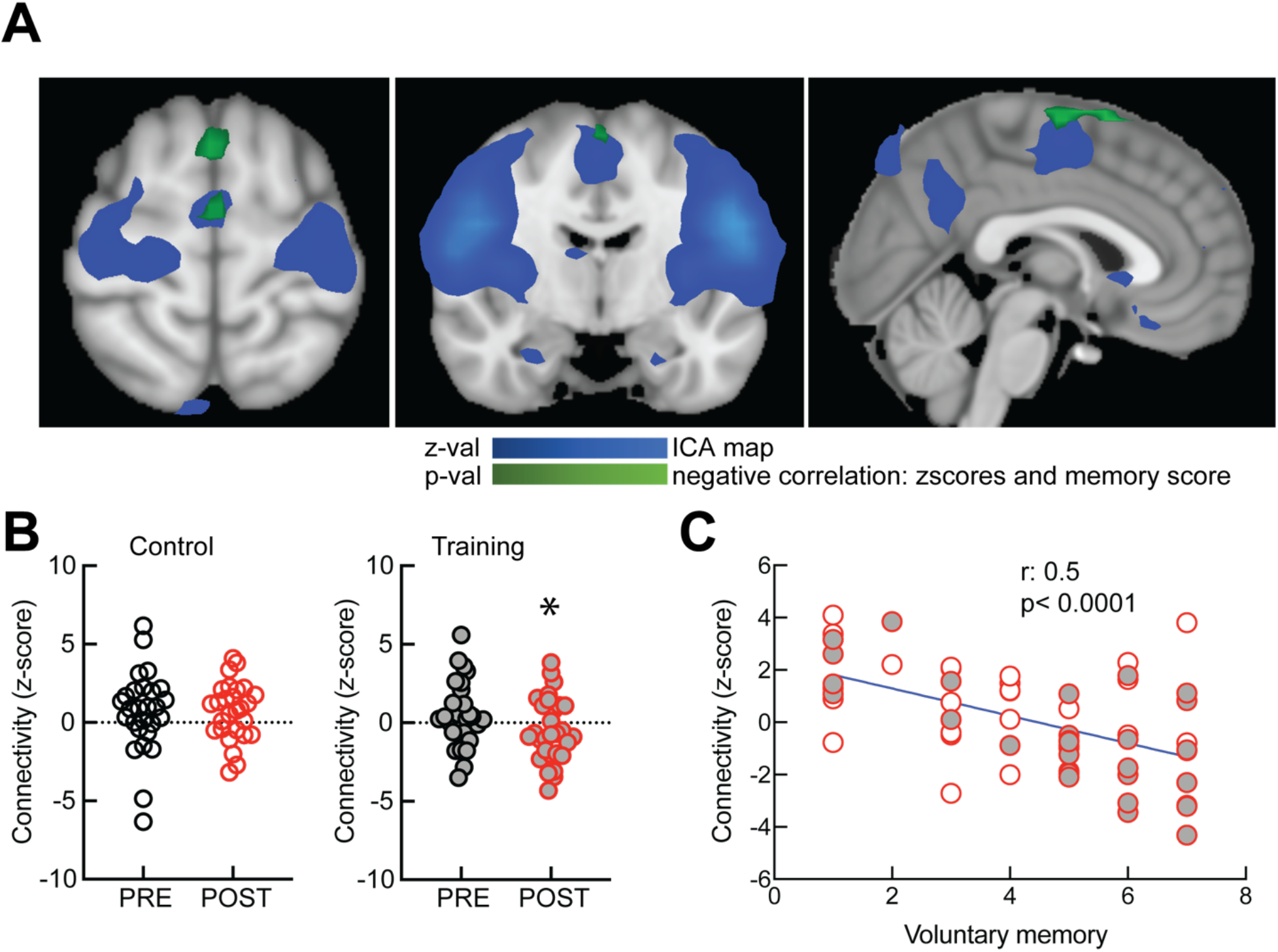
Sensorimotor network connectivity and improved voluntary memory after training. **A**. Clusters (green) within the sensorimotor network (blue) that negatively correlate with voluntary memory score. **B**. Subject specific z-score connectivity within the sensorimotor network in the training group (p=0.01) and the control group (p=0.9). **C**. Invers (negative) correlation between sensorimotor connectivity and voluntary memory scores across both training and control groups. Gray filled symbols represent experimental subjects. *, p<0.05.

In previous work, it has been shown that exercising sensorimotor behaviors leads to decreased BOLD activity, which was interpreted as a possible facilitation of neural functions, with more proficient sensorimotor skills associated with lower representation in sensory and motor structures (Kelly and Garavan, 2005). Consistent with these previous studies, we now find that exercising mental sensorimotor simulations of autobiographical memories leads to a decrease in intra-network connectivity that facilitates voluntary recall, as connectivity in the sensorimotor network negatively correlates with scores of voluntary memory for both experimental and control groups (**Fig. 3C;** r:0.5, p<0.0001).

### Immunological correlates of autobiographical training

Previous studies documented the role played by certain immunological factors, primarily TNFα, in memory and other cognitive processes (Besedovsky and del Rey, 2011; Liu et al., 2017; Morimoto and Nakajima, 2019). We investigated whether such biological processes could be implicated in autobiographical training (**Fig. 4**). We found that the levels of blood TNFα decrease after training (**Fig. 4B**, TNFα PRE: 355.1±60.54 pg/ml, TNFα POST: 199.1±43.03 pg/ml, Wilcoxon matched-pairs signed rank test, p=0.01), but not after the control procedure (TNFα PRE: 328.4±85.3 pg/ml, TNFα POST: 238.9±54.04 pg/ml, p=0.6). We performed a voxel-wise correlation with TNFα values, for subjects in POST-training condition. The TNFα values positively correlated with connectivity for clusters located in left motor area (**Fig. 4A**). We extracted these clusters and went back to subjects-specific connectivity maps to collect connectivity z-scores for both experimental and control groups in PRE and POST conditions. The level of circulating TNFα was positively correlated with this subject specific connectivity in the sensorimotor network (**Fig. 4C**; r:0.5, p=0.002). These findings could indicate a potential relationship between the decrease in blood TNFα and the neural correlates for improved voluntary recall following autobiographical training.

**Figure 4:**
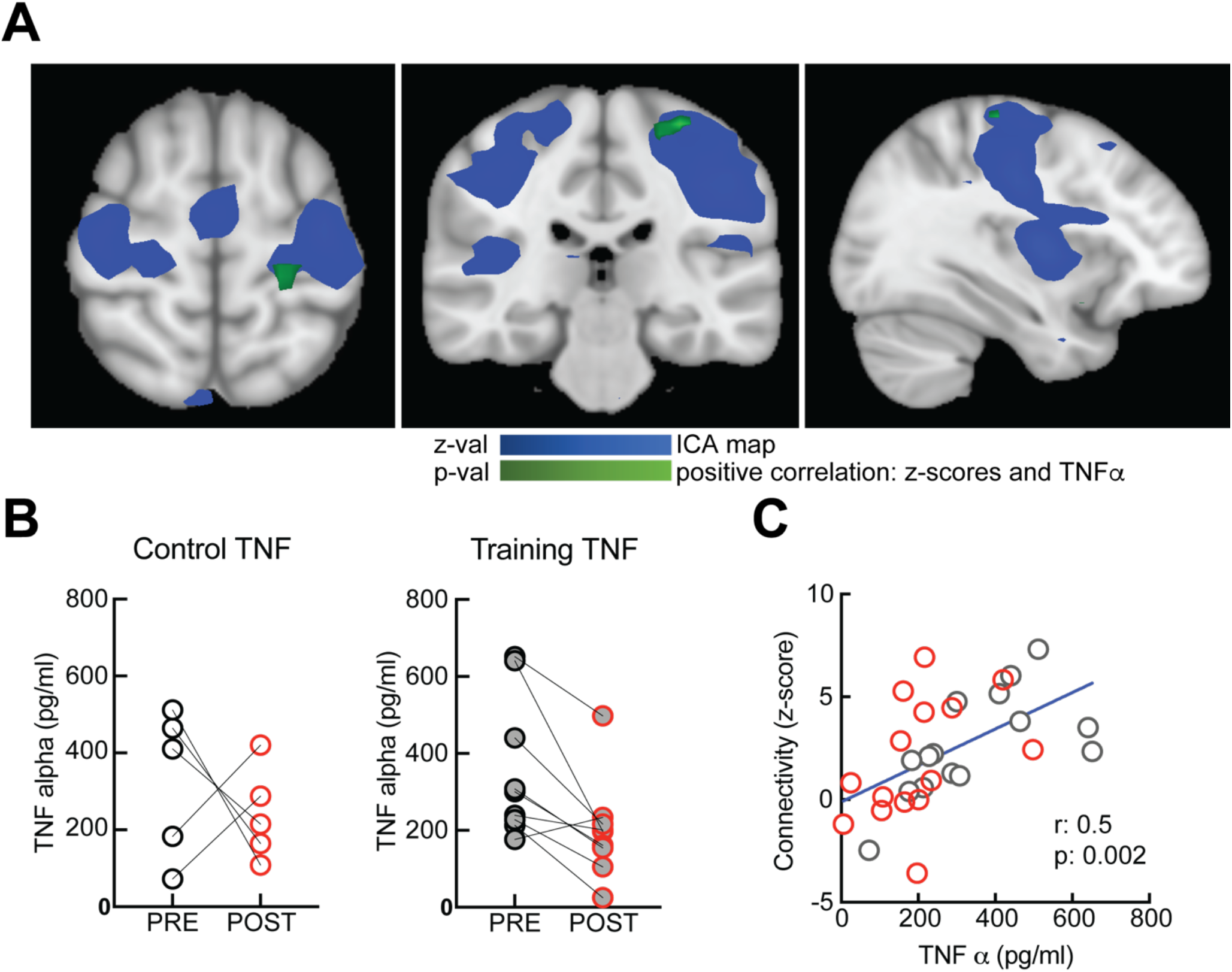
Correlation between TNFα and connectivity in the sensorimotor network. (A) Horizontal, coronal and sagittal sections showing the part of motor cortex (green) for which the connectivity with the rest of the sensorimotor network (blue) correlates with TNFα levels. (B) Summary data showing decreased TNFα levels after training (p=0.01) but not after control intervention (p=0.6). (C) Positive correlation between resting-state connectivity in the sensorimotor network and TNFα levels. Red symbols, training; gray symbols, control. *, p<0.05.

## Discussion

In this study we set out to determine if autobiographical training can improve mnemonic functions by changing connectivity in functional neural networks. We found that autobiographical memory training increased the number of odor-evoked retrieved memories. This confirmed previous studies in older subjects, showing that retrieval practice enhanced successful recall of personal events (Xu et al., 2020). We also found that autobiographical memory training increased the ease of voluntarily accessing memories outside the lab., consistent with previous clinical studies in schizophrenic patients (Ricarte et al., 2012).

A priori, we expected to find changes in the DMN network that would support the *scene construction* theory of memory, and in the sensorimotor network that would support the *embodied cognition* theory of memory. We found that autobiographical training leads to changes in the connectivity of both of these networks, which argues that both mnemonic theories have physiological relevance. None of the other thirteen functional networks considered showed significant changes with training.

In our study, autobiographical training increases connectivity between thalamus and DMN, and decreases connectivity within the sensorimotor network. Both of these changes could increase recall efficiency. Other studies using different training procedures found similar changes in DMN connectivity. Experienced mindfulness meditators (with more that 1000 hours of training) compared to beginner meditators (1 week of training) had increased connectivity between anterior DMN regions (dorso-medial PFC) and posterior DMN regions (inferior parietal lobule), suppoting the hypothesis that meditation training leads to functional connectivity changes between these two DMN hubs (Taylor, et al., 2013). An extensive study reviewing imagistic data on practice-related brain changes found that, as performance improves, a “process switch” allows for more efficient processing: as connectivity in specific brain circuits reorganizes metabolically costly brain activity decreases (Kelly and Garavan; 2004). This phenomenon is reminiscent of the synaptic plasticity of neural circuits described in animal models (Carcea and Froemke, 2013). The increased connectivity between right mediodorsal thalamus and the anterior DMN network found in our study could indicate that training enhances thalamic engagement in the recall process. In animal models, the mediodorsal thalamus has been identified as an important component of memory systems (Hsiao et al., 2020), and its main role could be to amplify and sustain representations in prefrontal structures (Schmitt et al., 2017; Parnaudeau et al., 2018). It is also possible that the increased connectivity that we detect after training reflects a stronger filtering input form prefrontal structures onto the thalamus (Nakajima et al., 2019), a process that would limit the interference of external sensory stimuli on the autobiographic recall.

The training-induced decrease in connectivity that we observe within the sensorimotor network is consistent with the ‘neural efficiency’ hypothesis that posits that training a response reduces activity in sensorimotor areas (Guo et al., 2017). In addition to decreases in activity, previous reports also found decreased connectivity following various types of training (McGregor and Gribble, 2017; Yue et al., 2020). However, to the best of our knowledge we are first to report decreased sensorimotor connectivity following autobiographical training. The invers correlation between sensorimotor connectivity and voluntary memory improvement post-training, supports the notion that this change represents a mechanism for ‘neural efficiency’.

The decrease in circulating immune factor TNFα indicates that autobiographical training might exert effects on bodily tissues as well. Immune response can be adjusted by the activity of the sympathetic nervous system and by hormonal activity especially linked to the adrenal gland (Segerstrom and Miller, 2004; Nance and Sanders, 2007; Kenney and Ganta, 2014). Two broad networks in the cerebral cortex that have access to adrenal gland (Dum, et al., 2016). The larger network they found includes all of the cortical motor areas in the frontal lobe and portions of somatosensory cortex, indicating that specific circuits exist to connect movement, cognition, and emotions to the function of the adrenal medulla. A systematic review of 24 functional magnetic resonance imaging studies investigated brain regions and networks associated with peripheral inflammation in humans and found a so-called “posterior putamen loop” which comprises also the sensorimotor cortex and is implicated in sensorimotor processes (Kraynak, et al., 2018). Taken together, these results indicate possible bidirectional interactions between peripheral inflammatory processes and various cognitive, affective and sensorimotor contexts.

A recent study suggests that mental health and physical health are linked by neural systems that regulate both somatic physiology and high-level cognition (Koban et al., 2021). The study proposes a “self-in-context” model which hypothesizes that events with personal meaning guide learning from experience and constructs narratives about the self and the environment (autobiographical memories), but at the same time can control peripheral physiology in a predictive way, including autonomic, neuroendocrine and immune functions. This model is in line with our findings and with previous research which demonstrated that cortical areas involved in the control of movement, cognition, and affect are sources of central commands to influence sympathetic arousal (Dum et al., 2016). This means that cognitive operations like action planning but also recalling significant actions from past events may be linked to the regulation of the adrenal function, and of the immune system. Finally, the Embodied Predictive Interoception Coding (EPIC) model proposed that brain did not evolve for rationality but to ensure resources for physiological systems within an animal’s body, and the psychological processes such as remembering, deciding and paying attention are in service for surviving, thriving and reproducing. To succeed, the brain has to control metabolic and other biological resources and performs by regulating the autonomic nervous system, the endocrine system and immune system (Kleckner et al., 2017).

In the future it will be important to determine the mechanism by which autobiographical training could impact the levels of TNFα. Given the correlation between TNFα levels and sensorimotor connectivity, structures within the sensorimotor network could be part of the mechanism for autobiographical immune control. Limitations of this study are represented by the little success in recruiting male subjects into the study, and by the small sample of reliable immunitary data collected from the participants.

## Funding

The project “Developing a methodology of therapy through theatre with an effect at the neurochemical and neurocognitive levels” (MET) is co-financed by the European Regional Development Fund (ERDF) through Competitiveness Operational Programme 2014-2020, SMIS code 106688 and implemented by UNATC “I.L. Caragiale”, CINETic Centre, LDCAPEI LAB. First author was also funded by an International Brain Research Organization (IBRO) fellowship.

## Disclosure

The authors have nothing to disclose. The authors declare that the research was conducted in the absence of any commercial or financial relationships that could be construed as a potential conflict of interest.

## Acknowledgements

We thank Prof. Nicolae Mandea, Prof. Liviu Lucaci and Prof. Radu Apostol for their administrative support, Dr. Robert C. Froemke and Dr. Justin S. Riceberg for consultation, and Doina Strat for her technical support.

## Author contributions

All authors contributed to the design of experiments and interpretation of results. DC (first author) performed the autobiographical training with help from AS, TR, M-YT, AD, and collected behavioral and MRI data with help from DG, DT and AIB. MO collected and analyzed the MRI data. CCP analyzed the behavior data with help from IRP. NP recruited and selected subjects. DC, MO and IC wrote the manuscript, with feedback from all authors.

## Notes

### Competing Interest Statement

The authors have declared no competing interest.

